# Binocular System with Asymmetric Eyes

**DOI:** 10.1101/241323

**Authors:** Jacek Turski

**Affiliations:** Department of Mathematics and Statistics, University of Houston-Downtown, One Main Street, Houston, TX 77002

## Abstract

I elaborate binocular geometry with a novel eye model that incorporates the fovea’s temporalward displacement and the cornea and the lens’ misalignment. The formulated binocular correspondence results in longitudinal horopters that are conic sections resembling empirical horopters. When the eye model’s asymmetry parameters’ range is that which is observed in healthy eyes, abathic distance also falls within its experimentally observed range. This range in abathic distance is similar to that of the vergence resting position distance. Further, the conic’s orientation is specified by the eyes’ version angle, integrating binocular geometry with eye movement. This integration presents the possibility for modeling 3D perceptual stability during physiological eye movements.

**OCIS codes:** (330.1400) Vision - binocular and stereopsis; (330.4060) Vision modeling

## 1. INTRODUCTION

In the human visual system, the left and right retinae receive slightly different projections of a scene due to eyes being forward-facing and horizontally separated from one another. Despite the disparity in retinal projections, we perceive the world as if it were seen from just one viewpoint, the brain taking advantage of this two-dimensional disparity to provide our perception with a vivid impression of depth, or stereopsis. Two basic concepts useful to understanding these phenomena are the horopter and the cyclopean axis. Here I investigate these concepts with binocular geometry and a novel eye model.

### A. Horopter: Basic Concepts

This review aims to give a short background that includes basic horopter models and their discriminating geometric facts. Please, refer to [1] for a comprehensive survey of stereoscopic vision and to [2] for an account of the long history underpinning our notion of the horopter in binocular theory of vision.

When a point source of light, such as a star, is in the binocular field of vision, it is projected into the two eyes along different lines where it stimulates light-sensitive cells in the retinae. Two retinal elements, i.e., receptive fields [3], one in each eye of the binocular system, are called corresponding points if they invoke a single perceptual direction for the spatial point stimulating them. The foveal centers are corresponding points stimulated by the fixated point whose perceived direction is called either the principal visual direction or the cyclopean axis. The visual directions of all other points in the visual field are perceived in relation to this principal direction.

The binocular correspondence problem is the delineation of the horopter defined as the locus of all spatial points stimulating corresponding points. But during simultaneous stimulation of both retinae, the binocular correspondence cannot be localized with any greater precision than the area where retinal images are fused as a single percept. The range of the spatial region that the fusion occurs within is known as Panum’s fusional area. However, when the two retinae are stimulated separately in the procedure known as the nonius method, a more precise binocular correspondence can be established [4].

The analytical expressions of the longitudinal horopter curves can be obtained using an eye model, or, alternatively, by curve-fitting data collected in experiments delineating binocular correspondence. However, because eye models simplify binocular geometry differently, each provides a distinct analytic form of the horopter curve. Also, depending on the experimental method used to delineate binocular correspondence [5], different curves may be used to fit data obtained from different experimental paradigms.

The Vieth-Müller circle (VMC) is commonly referred to as the longitudinal horopter derived from geometric analysis of binocular projection. However, its construction relied on an eye model that assumes, incorectly, that the nodal point coincides with the eye’s rotation center [6]. Under this assumption, each VMC passes through the fixation point and connects the rotation centers of the two eyes.

Recently, I constructed a family of geometric horopters in [7] by correcting the nodal point’s location as required by the eye’s anatomy to almost midway between the eye’s rotation center and pupil. Each longitudinal horopter curve is then a circle that passes through the fixation point and connects the eyes’ nodal points. When the fixation point moves along the VMC, the eyes’ rotation centers are fixed and the VMC remains unchanged. But because the nodal points’ positions change when the fixation point changes, the longitudinal horopter curves in [7] do change as the fixation point moves. More analysis of the difference between the VMC and the geometric horopter in [7] is given in the last section.

Nevertheless, two anatomically incorrect assumptions were used in each of the horopter models discussed above. First, the fovea centralis was located at the posterior pole of the eyeball. Second, the nodal points’ and optical axes’ congruence in the two eyes of the binocular system brought the geometrically defined corresponding points into congruence.

In the human binocular system, the fovea is displaced temporalward from the posterior pole. Further, the empirical horopter is flatter at the fixation point than the circular horopter. This difference between empirical horopters and geometric horopters, or the Hering-Hillebrand horopter deviation, is depicted in Figure 1 for symmetrically converging eyes.

**Fig. 1.**
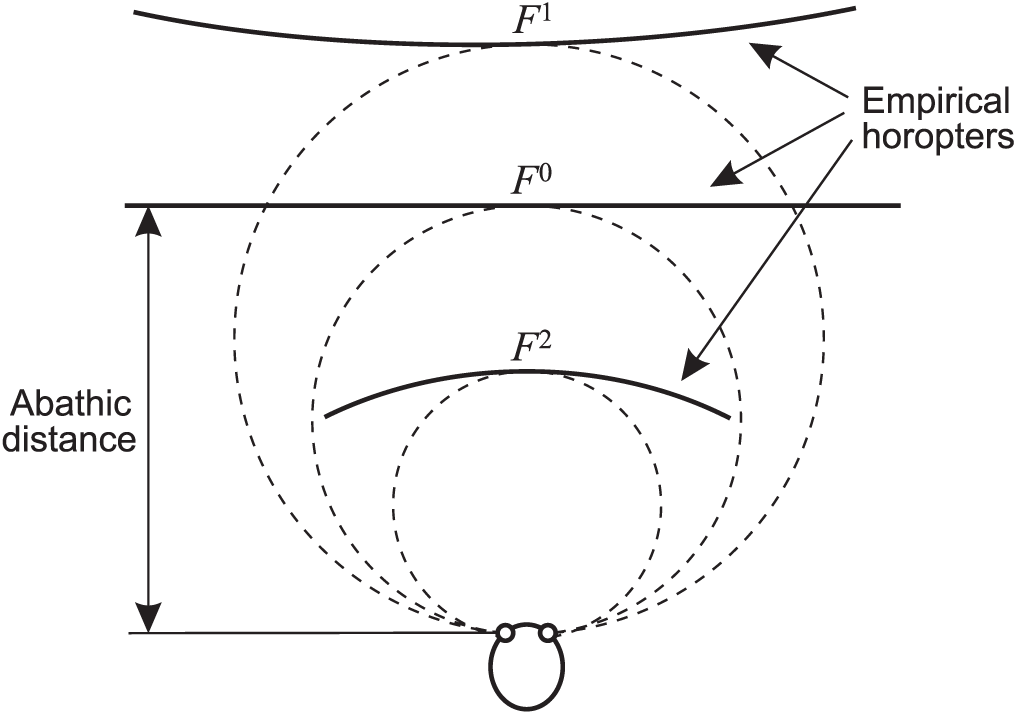
Empirical longitudinal horopters shown schematically for fixation points *F*^0^, *F*^1^ and *F*^2^. Each horopter near the fixation point determines the perceived frontal line. For the fixation point *F*^0^, the apparent frontal line agrees with the objective frontal line. The VMCs are shown in dashed-lines.

We see that the shape of each horopter curve near the fixation point changes: for the fixation point *F*^0^ at the abathic distance, the horopter curve is a straight line and the curvature is zero, for forward fixations with larger distances than the abathic distance, the curvature is negative, and for such fixations with shorter distances, the curvature is positive. The fact that the empirical horopter changes shape with fixation distance shows that the corresponding points cannot be congruent in both eyes. Please, refer to [8, pp. 49–52] where the necessary conditions for the congruence of the corresponding points are discussed.

### B. The Paper Contributions

In this paper, I study the geometric properties of the longitudinal horopters in a binocular system with a novel eye model that corrects the anatomically inaccurate assumption made in [7] that located the fovea centralis at the posterior pole. This model incorporates the fovea that is displaced temporalward from the posterior pole and uses a single refractive surface to represent the cornea and the crystalline lens’ misalignment observed in human eyes.

I refer to this anatomically extended reduced eye as the asymmetric eye. One immediate consequence of using the asymmetric eye in a binocular system is that the corresponding points are not congruent in the retinae.

Each horopter point in the horizontal visual plane is derived by finding the intersection of the visual lines that are passing through the different retinal points of one pair of corresponding points. For each eye’s fixation, the curve passing through these points turns out to be a conic section resembling the empirical horopter, its geometry fully specified by the eye model’s asymmetry. In particular, values of the refractive surface’s asymmetry parameter in the range similar to that of the crystalline lens’ tilt, as measured in healthy eyes, also produce values of abathic distance that are in the range observed in human vision. Also, the version angle of each eyes fixation is equal to the angle that specifies the orientation of the conic section describing the longitudinal horopter. This equality integrates eye movement with binocular geometry.

This study provides anatomical support to the only other comprehensive modeling of empirical horopters as conic sections, carried out in [9] and [10] by methods of analytic geometry. The equations introduced on an ad hoc basis by Ogle in [9] for the forward gaze and extended by Amigo in [10] to any horizontal gaze were used to obtain a family of conics whose free parameters had to be determined experimentally for each subject. Later I will demonstrate the numerical similarities and geometric differences between the horopter curves obtained in Ogle and Amigo’s classical work and those obtained here.

Finally, I will discuss the two research problems that follow naturally from my study. The first deals with the potential relationship between the eye’s fixation at the abathic distance and the vergence resting position during degraded visual conditions. The second problem involves the eye’s movement and binocular geometry integration that is needed in robotic modeling of the visual and oculomotor processes that maintain a stable 3D perception during saccadic and smooth pursuit eye movements.

Preliminary results of this work were presented in [11].

## 2. RELATED WORK BY OGLE AND AMIGO

I review here Ogle and Amigo’s classical results, in doing this I follow [12] and [10]. Later, I will compare their results with those I obtain in this paper.

For symmetrically convergent eyes, horopter curves are conic sections determined from the set of numbers *α_l_* and *α_r_* that satisfy the family of equations

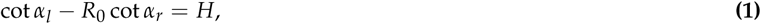

introduced by Ogle in 1932. In (1), *H* and *R*_0_ are nonnegative parameters. Also, *α_l_* and *α_r_* are the angles subtended at the rotation centers of the left and the right eyes by *F*, the fixation point, and *P*, any other point on the longitudinal horopter; see Figure 2.

**Fig. 2.**
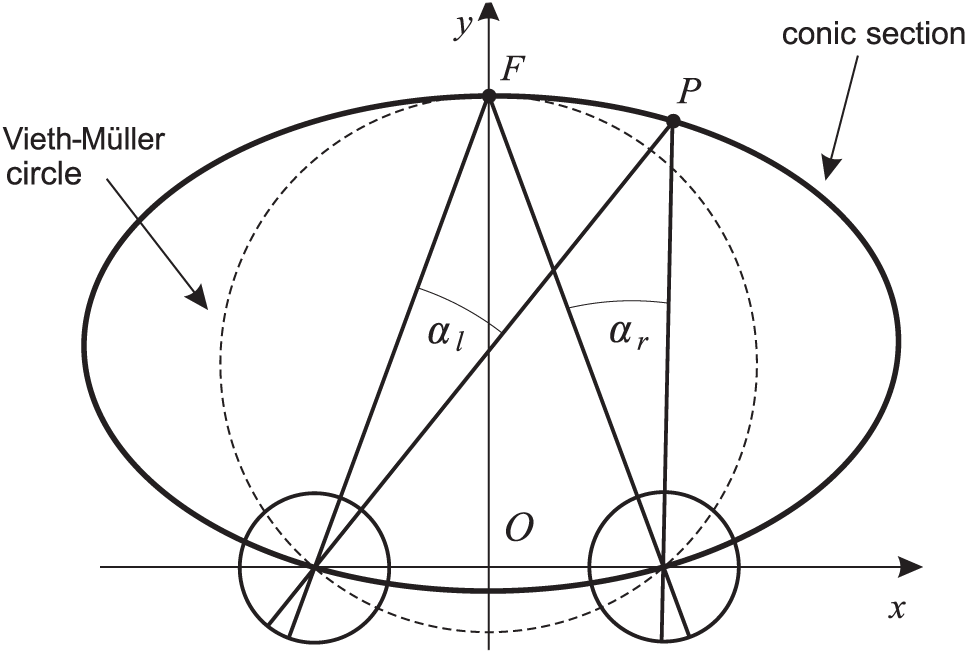
The conic section for the fixation point F determined from Ogle’s condition (1) with *R*_0_ = 1.

The parameter *H* characterizes the curvature of the horopter near the fixation point. The parameter *R*_0_ accounts for the amount by which the horopter curve is skewed at the symmetrically fixated point. When *R*_0_ = 1, the horopter curves are symmetrical around the fixation point, cf. Figure 2.

Ogle justified (1) by fitting the equation to data obtained in experiments measuring longitudinal horopters. He estimated from the plot of the equation that the values of *R*_0_ are very close to unity and the values of *H* are less than o.2.

In this study, I assume *R*_0_ = 1 in Ogle’s family of horopter-defining equations

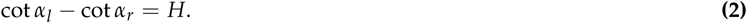

From this equation, by expressing *α_l_* and *α_r_* in coordinates (*x*, *y*), as shown in Figure 2, Ogle derived the following family of conic sections:

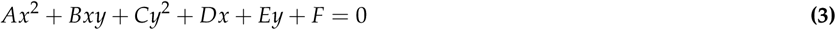

with the coefficients,

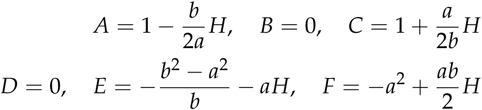

where 2*a* is the interocular distance and *b* is the distance from the origin *O* of the coordinate system to the point of symmetric fixation. When

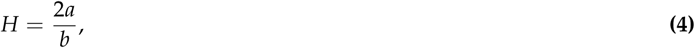

the conic section is a straight line *y* = *b* passing through the fixation point. Thus, in this case, the apparent frontal line agrees with the objective frontal line. Further, if 2*a/b* > *H* > 0, the curve is an ellipse, and if *H* > 2*α/b*, the curve is the hyperbola’s branch passing through the fixation point. Moreover, if *H* ≠ 0, these conic equations are reduced to the VMCs. The parameter *H* ≠ *0* describes the Hering—Hillebrand deviation.

The results of Ogle were extended by Amigo in 1965, who obtained conic sections for when the eyes are in asymmetric convergence. His conics are given in the coordinate system (*X*, *Y*), as shown in Figure 3. Thus, the coordinate system used by Ogle is one rotated by the angle γ′ with respect to the coordinate system used by Amigo. As usual, the counter-clockwise angles are taken to be positive, hence γ′ *>* 0. However, I use the angle γ = − γ′ to emphasize geometric transformations that are applied to points, rather than that which are applied to the coordinate system. We note that *F*(0, *b*) expressed in (*X*, *Y*) coordinates takes on the form *F*(−*b* sin *γ*, *b* cos *γ*) when expressed in (*x*, *y*)-coordinates.

**Fig. 3.**
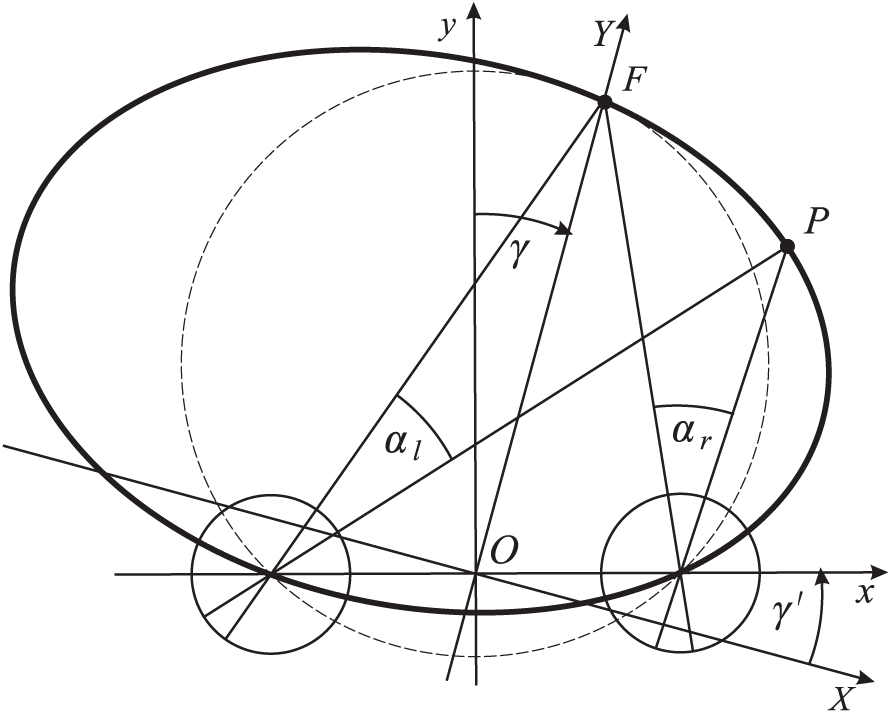
The coordinated (*X*, *Y*) used by Amigo in deriving the family of conic sections. One conic section for the fixation point *F* is shown by a solid line.

The Amigo family of conic curves given in the equation (5) in [10], is transformed to the (*x*, *y*)-coordinate system used by Ogle to unify the description of the horopter curves. I do this with the help of the transformation

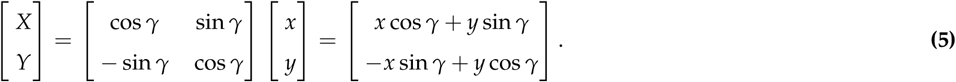

Thus, I change *x* to *x* cos *γ* + *y* sin *γ* and *y* to −*x* sin *γ* + *y* cos *γ* in the equation (5) in [10] to obtain the following expressions:

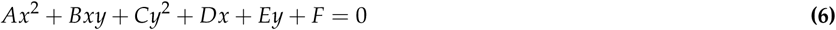

with the coefficients

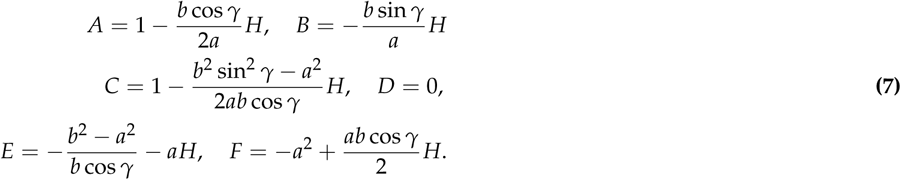

We note that *D* = 0 means that the VMC obtained from (6) with *H* = 0 has its center on the *y*-axis at (0, (*b*^2^ − *a*^2^)/2*b* cos *γ*) and its radius as *r* = ((*b*^2^ − *a*^2^)^2^/4*b*^2^ cos^2^ *γ* + *a*. The slope of the tangent line to the conic at the fixation *F*(−*b* sin *γ*, *b* cos *γ*) is obtained by the implicit differentiation of (6) at *F*,

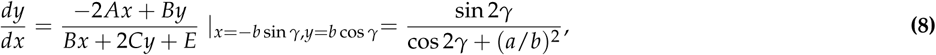

giving this equation for the tangent line at *F*,

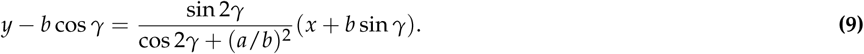

Because this slope does not depend on *H*, the tangent to the VMC (for *H* = 0) at *F*(−*b* sin *γ*, *b* cos *γ*) is the same as in equation (9). Further, following Amigo [10], upon taking

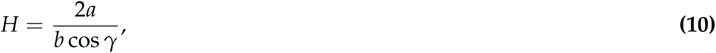

equation (6) reduces into two equations, *y* = 0 and (9). When *γ* = 0, (10) reverts to (4), reducing the conics (3) into the straight line *y* = *b* at the abathic distance *b*. The straight lines associated with (10) represent conics when the discriminant associated with (6) vanishes, cf. Amigo [10]. Line (9) gives the apparent frontal line that coincides with the objective frontal line. Amigo called (10) the representation of the general expression for abathic distance.

The orientation of a general conic in (6) was, surprisingly, not considered by Amigo, but can be given by the angle

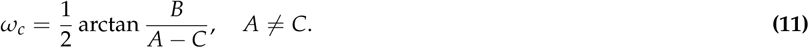

This is the angle of rotation that transforms this conic into its standard orientation, which is when conic axes are parallel to coordinate axes. Introducing the coefficients (8) into (11), we obtain

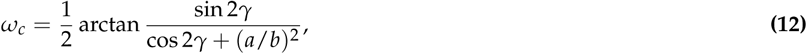

which is independent of the parameter *H*.

There have been other attempts to extend Ogle’;s original work [13–15]. However, these attempts are either less comprehensive or less amenable to analytical treatments than those performed by Amigo. Additionally, there have been other approaches to modeling horopter curves. Most notable of these is the study of the geometric structure of visual space carried out by Luneburg [16], who developed the model of a metric visual space of constant hyperbolic curvature. He identified the horopter curves with the geodesic lines of his metric and pointed out their resemblance to empirical longitudinal horopters when the eyes are symmetrically fixated on points in the horizontal plane.

Please refer to [17] for a review of the intricate relationship between geometry and spatial vision.

## 3. THE ASYMMETRIC EYE IN BINOCULAR SYSTEM

### A. The Eye Optical System

The eye’;s optical system comprises of four main optical components: the cornea, lens, pupil and a light-sensitive retina. The pupil’s center and the refractive surfaces curvature centers of the cornea and the lens are not perfectly aligned. Although, strictly speaking, an optical axis that would pass through the curvature centers does not exist, a reference line defined as the best fit through these points can well approximate an optical axis for many applications [18]. Even so, the fovea centralis is displaced 1 to 2 mm temporalward from the intersection of the optical axis with the retina.

Although the fovea’s displacement contributes to optical aberrations that degrade the retinal image, the eye’s sophisticated aplanatic design compensates for some of the resultant limitations in optical quality with its nonaxial geometry of refractive surfaces and the distribution of the gradient refractive index of the lens [19].

### B. The Asymmetric Eye

Eye models range from accurate optical models aiming to comprehensively represent ocular anatomy and physiology, to simpler models that, while not usually anatomically accurate, are developed to mimic only specific eye functions. For a review of a selection of commonly used eye models and a discussion of their advantages and disadvantages, refer to [18].

The law of parsimony suggests that the best functional models will include only those details of the eye absolutely necessary for accomplishing the model’s purpose [18]. Following this law, the eye model constructed here incorporates the most important features of the human eye’s asymmetric design: the fovea’s displacement on the retina from the posterior pole and the cornea and crystalline lens’ misalignment [20, 21].

This asymmetric eye, which minimally extends the reduced eye in Figure 4 (a), is schematically shown in Figure 4 (b) for the right eye. The optical axis is defined here as the line passing through the nodal point *N* and the eye’s rotation center *C*. The misaligned cornea and lens are represented by a single refractive surface that is tilted nasally relative to the optical axis by the angle *β* at the nodal point. This tilt is accounted for in the binocular analysis by representing the refractive surface with the image plane obtained by rotating, by the angle *β* at the eye’s rotation center *C*, the plane that intersects, perpendicularly, the optical axis at *C*, cf. Figure 4 (b). This image plane allows us to use, in the next section, a simple formulation of the binocular correspondence.

**Fig. 4.**
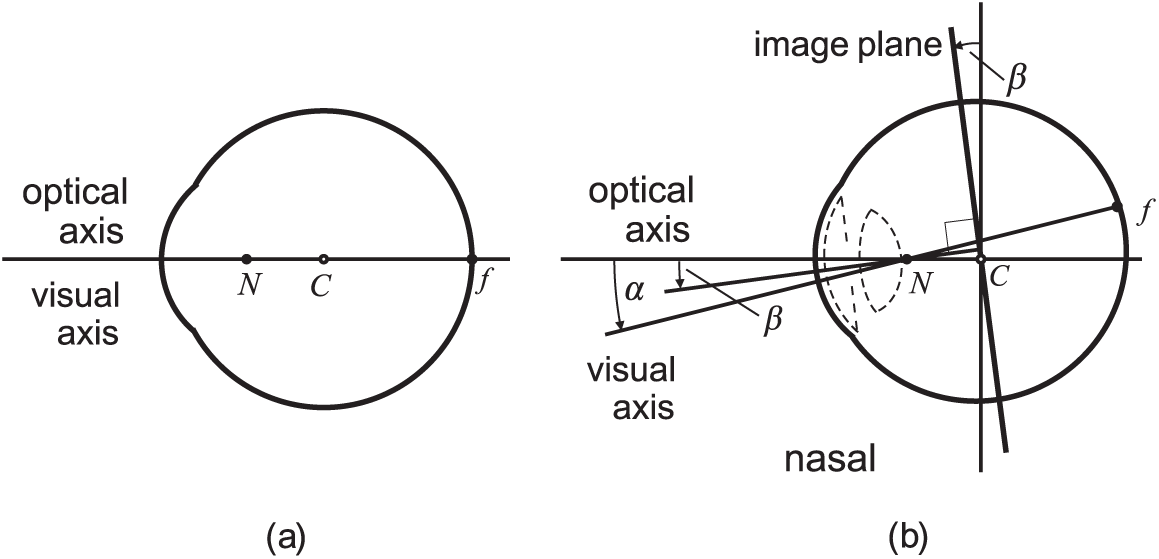
(a) The reduced eye. The anterior pole, the nodal point, *N*, the eye rotation center, *C*, and the fovea centralis, *f*, are all located on the optical axis. The nodal point is 0.6 cm anterior to the rotation center. (b) The asymmetric eye, shown here for the right eye, minimally extends the reduced eye. The cornea and lens are not included in the asymmetric eye.

The refractive surface tilt introduces its decentration *e* relative to the optical axis in the nasal direction

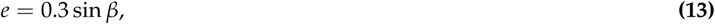

where 0.3 cm is the assumed distance from the nodal point to the point at the middle of the lens on its optical axis. In both eye models in Figure 4, the separation of the nodal point from the eye’s rotation center is 0.6 cm.

The visual axis connects the fixation point with the fovea centralis, *f*, and passes through the nodal point *N*. Because the fovea centralis is displaced 1 to 2 mm temporalward, the visual axis is angled nasally from the optical axis by the angle *α* with its values ranging from 4 to 7 degrees. In the asymmetric eye, I assume the average value *α* = 5.2°.

I should mention that different axes and angles have been used in measurements of the eye’s misaligned optical components [20, 21], which are later compared with those used in the asymmetric eye. However, the angle *α* has the least amount of inter-patient variability and is the most reliable benchmark for refractive surgery [22].

### C. The Binocular Correspondence

In the binocular system with asymmetric eyes, points in space are projected through the nodal point *N* of each eye into both the sphere representing the retina and the image plane.

For a given fixation point, the left eye’s visual axis passes through the respective nodal point and intersects the retina at the fovea’s center, *f_l_*, and the image plane at point *O_l_*. The other visual axis passes, similarly, through the nodal point of the right eye before intersecting the retina at the fovea’s center and the image plane at *O_r_*.

The corresponding points in the binocular system with the asymmetric eye are defined as follows. First, I choose a pair of points, one in each of the eyes’ image planes, which are intersected by the horizontal visual plane. If these points are the same distance away from their respective points *O_l_* and *O_r_* and are also on the same side of these points, their projections through the nodal points to the spheres (i.e., retinae) will define the corresponding points. Sometimes, for the sake of simplicity, both the corresponding points on the retinae and the projection-related points on the image planes are referred to as corresponding points.

### D. The Abathic Distance

The asymmetric eye-position shown in Figure 5 is symmetrically convergent, with both image planes perpendicular to the *y*-axis and coplanar. I will prove below that this is the abathic distance position; i.e., that the horopter is a straight line for this position.

**Fig. 5.**
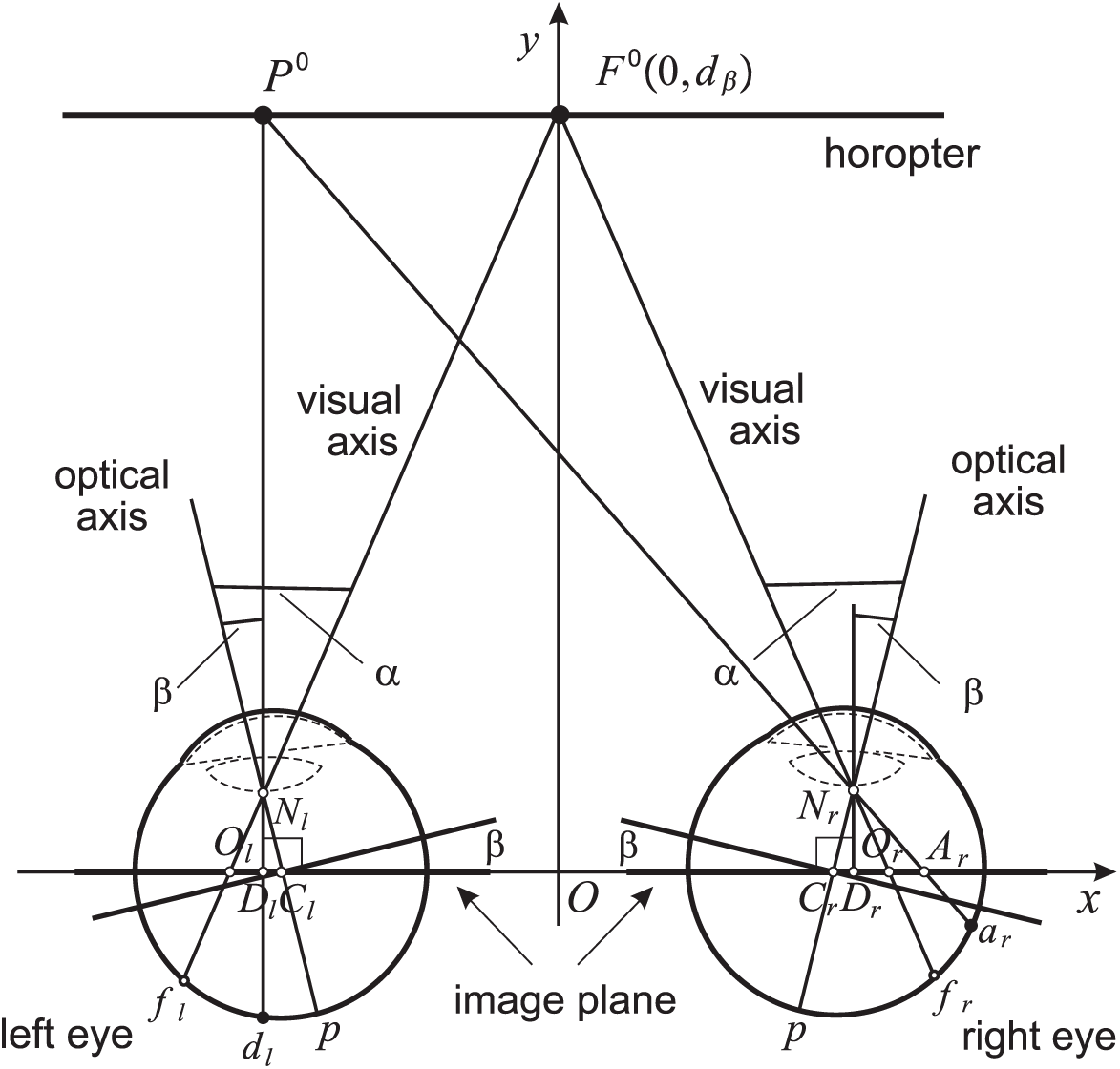
The binocular system with the asymmetric eye shown in the symmetric convergence at the abathic distance *d_β_*.

For a given *α*, *β*, the ocular separation 2*a* = |*C_l_C_r_*| and |*C_l_ N_l_* | = 0.6, the abathic distance can be obtained from two similar triangles, △*O_l_OF*^0^ and △*O_l_D_l_N_l_*, as follows

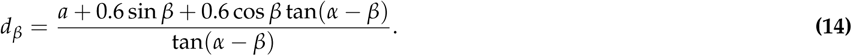

In (14), I also used that |*C_l_D_l_* | = 0.6 sin *β* and |*D_l_O_l_* | = 0.6 cos *β* tan(*α* − *β*), which can be obtained from the right triangles △*C_l_D_l_ N_l_* and △*O_l_D_l_ N_l_*, respectively.

Further, by using similar triangles △*D_l_P*^0^ *A_r_* and △*D_r_N_r_ A_r_*, we see that, for the corresponding points *D_l_* and *A_r_* in the image plane, |*D_l_P*^0^| = *d_β_*, where *d_β_* is given in (14). This proves that the horopter through *F*^0^ is indeed the straight line shown in Figure 5.

The retinal corresponding points *d_l_* and *a_r_* given by the choice of points *D_l_* and *A_r_* in the image plane are of different distances from their foveae, as can be seen in Figure 5. Now, equally dividing the intervals *O_l_D_l_* and *O_r_ A_r_* as required in the construction of corresponding points, and projecting this division points into the retinae of the respective eyes, we see that the obtained corresponding points are compressed, in the temporal retinae, relative to those in the nasal retinae. We conclude that for the binocular system with asymmetric eyes, the conditions for the corresponding points’ congruence discussed in Section 1 A cannot be satisfied.

Using the average value of 2*a* = 6.5 cm, the angles *α* = 5.2° and −0.4°≤ *β* ≤ 4.7° in (13) and (14), we find the lens’ values of the nasal decentration relative to the optical axis in the range −0.02 mm ≤ *e* ≤ 0.25 mm and values of the abathic distance in the range

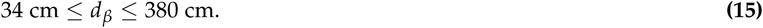

The lens’ tilt and decentration in human eyes are measured in different ways as the result of using different techniques. In [20], the tilt and decentration values were obtained relative to the corneal axis while, in [21], the tilt was obtained relative to the fixation axis and the decentration was obtained relative to the pupil center.

Although, the asymmetry parameters of the refractive surface are defined relative to the optical axis, the assumed values of the tilt given by angle *β* and the calculated decentration *e* are of the order of the corresponding values reported in [20, 21]. Moreover, the calculated range in (15) agrees with the observed values of the abathic distance in human vision [23, 24], its average value being

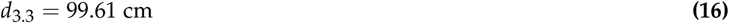

for the value *β* = 3.3° [25]. We also see that when the value of *β* increases to 5.2°, the abathic distance in (14) increases to ∞, and this has been occasionally reported in experimental measurements.

## 4. LONGITUDINAL HOROPTER CURVES

The notation used in the analytic derivation of the horopter’s points are presented in Figure 6. This figure only shows the directions of the visual lines in order to emphasize the corresponding points on the image planes that were used in calculations.

**Fig. 6.**
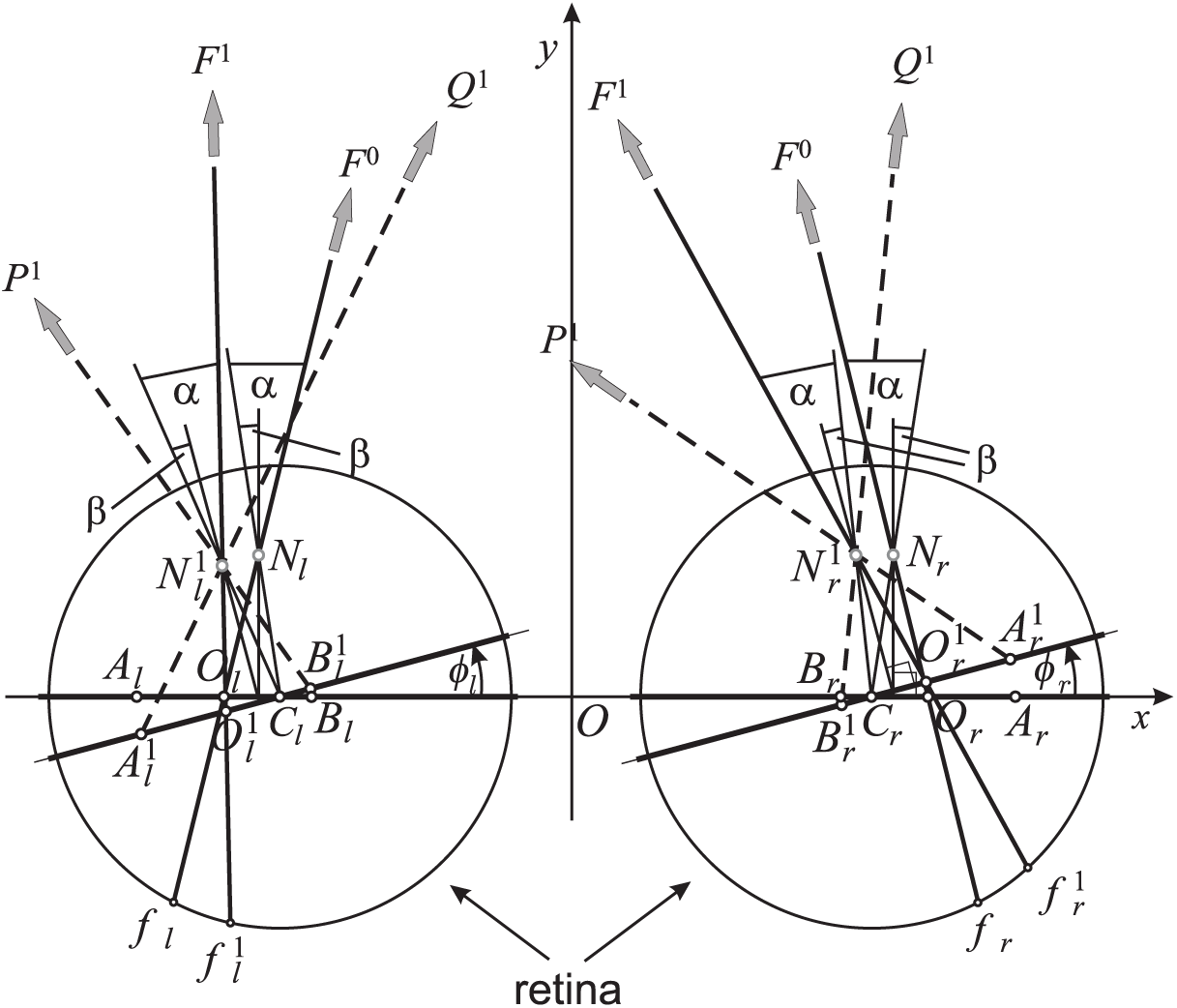
Geometric details and notation used in the text to derive the points on the longitudinal horopters specified by the eye’s asymmetry. *F*^0^ is the fixation point at the abathic distance. Only directions of the visual lines of the points on the horopter curves are shown.

The fixation point *F*^1^ is obtained by rotating eyes from the symmetric fixation *F*^0^ at the abathic distance *d*_3.3_ by the angles *φ_l_* and *φ_r_* for the left and right eye respectively. A pair of corresponding points are specified on the image plane by a distance *c* from the *O_l_* and *O_r_*. Figure 6 shows three pairs of corresponding points on each of the two image planes: (*A_r_*, *B_l_*), (*O_r_*, *O_l_*), (*A_l_*, *B_r_*) for the fixation point *F*^0^ and 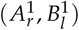, 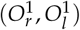, 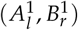 for the fixation point *F*^1^. Here, the letter ‘*A*’ indicates points on the temporal side and the letter ‘*B*’ indicates points on the nasal side of the respective image plane’s foveal centers. The visual lines’ directions (dashed lines) of horopter points *P*^1^ and *Q*^1^ that project to two pairs of corresponding points 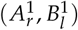 and 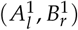 are shown only for the fixation *F*^1^.

I use vector algebra to calculate the horopter points. Using Figure 6, the position vector of the point 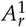 can be expressed in terms of binocular system parameters by its position vector,

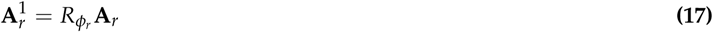

where

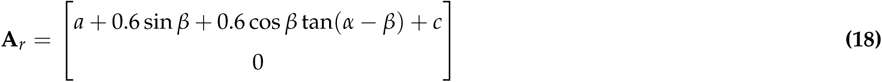

and 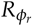 is the two-dimensional rotation matrix given by

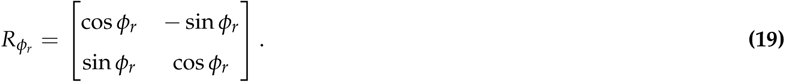

In the expression (18), I used the relations similar to those used in the derivation of (14), just obtained from the triangle △*C_r_N_r_O_r_*.

Further, the direction vector 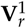 of the line passing through point 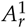 and the horopter point *P*^1^, the visual line of *P*^1^ when the eye is fixating on *F*^1^ can be written as

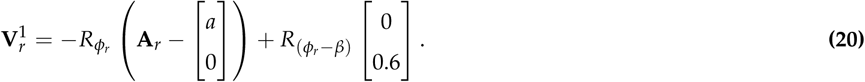

Similar expressions can be obtained for the left eye:

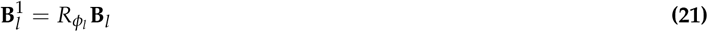

with

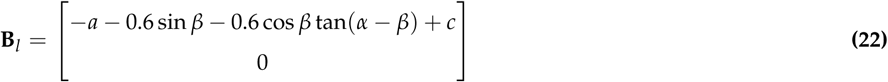

and

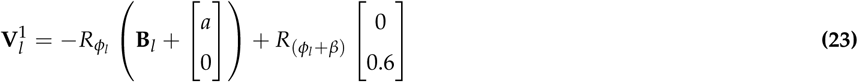

The horopter point *P*^1^ lies on the intersection of the two visual lines of corresponding points 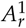 and 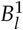. The coordinates of point *P*^1^ are derived by solving the vector equation

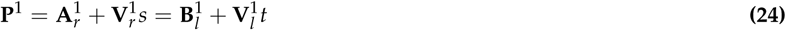

for (*s*, *t*).

Coordinates of the point *Q*^1^ can be obtained in a similar way. Also, if we take *c* = 0 in the above expressions, we get the description of the fixation point *F*^1^ specified by the values of *φ_l_* and *φ_r_*.

### A. Points Along Horopters

I proved in Section 3 that the horopter curve for the symmetrically fixated point *F*^0^(0, *d*_3_._3_) is a straight-line. Here, I will compute points on the two horopter curves, each for different asymmetric fixation obtained from *F*^0^(0, *d*_3_._3_). Fixation point *F*^1^ is obtained by rotating the eyes with the angles *φ_l_* = 10°, *φ_r_* = 9° and fixation point *F*^2^ is obtained by rotating the eyes with the angles *φ_l_* = −15°, *φ_r_* = −13°.

The computations of the horopter points are performed using the human binocular system’s average dimensions. Note that although the real numbers’ floating-point representations is rounded in my calculations to 10 decimal places, as required for geometric conclusions, they are shown here to only 3 significant places for simplicity’s sake.

The points on the horopter for the fixation *F*^k^, *k* = 1, 2, are denoted as follows: *P*^k^ and *Q*^k^ with *n* = 1 for *c* = 0.025 cm, *n* = 2 for *c* = 0.075 cm, *n* = 3 for *c* = 0.125 cm and *n* = 4 for *c* = 0.4 cm. The point *F*^k^ is obtained by taking *c* = 0.

Substituting *α* = 5.2°, *β* = 3.3°, *φ_l_* = 10°, *φ_r_* = 9° and *c* = 0.4 into the expressions from (17) to (23), which are then used in (24), we obtain

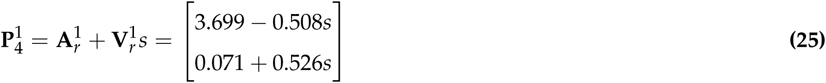

and

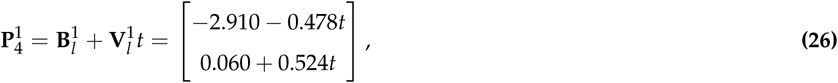

which can be written as the system of linear equations

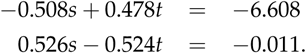

The solution of this system

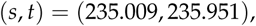

used, for example, in (25), gives coordinates of point *P*_14_:

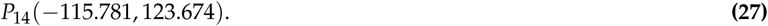

Similarly, for the coordinates of the point 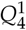:

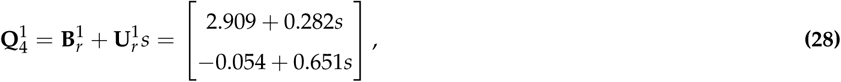

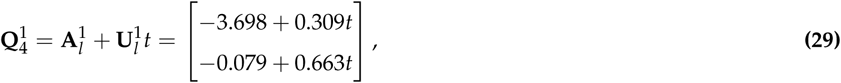

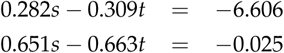

with the solution of this system

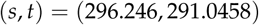

and

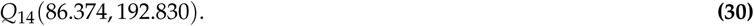

Finally, taking *c* = 0, I obtain the fixation point *F*^1^(−22.0276, 131.552).

The points computed on the horopter curves for fixations *F*^1^ and *F*^2^ are listed in Table 1.

**Table 1.** The Horopter Points for Fixations *F*^1^ and *F*^2^

| $F^1 : \phi_l = 10^\circ, \phi_r = 9^\circ$ | $F^2 : \phi_l = -15^\circ, \phi_r = -13^\circ$ |
| --- | --- |
| Horopter Points |  |
| $F^1(-22.028, 131.552)$ | $F^2(15.358, 61.434)$ |
| $P_1^1(-27.319, 129.821)$ | $P_1^2(12.931, 62.623)$ |
| $Q_1^1(-16.688, 133.458)$ | $Q_1^2(17.708, 60.141)$ |
| $P_2^1(-37.836, 126.863)$ | $P_2^2(7.919, 64.936)$ |
| $Q_2^1(-5.784, 137.828)$ | $Q_2^2(22.164, 57.395)$ |
| $P_3^1(-48.396, 124.550)$ | $P_3^2(2.7280, 66.919)$ |
| $Q_3^1(5.5630, 143.009)$ | $Q_3^2(26.244, 54.469)$ |
| $P_4^1(-115.781, 123.674)$ | $P_4^2(-25.598, 72.291)$ |
| $Q_4^1(86.374, 192.830)$ | $Q_4^2(40.601, 37.244)$ |
| Nodal Points |  |
| $N_l^1(-3.388, 0.584)$ | $N_l^2(-3.128, 0.588)$ |
| $N_r^1(3.190, 0.597)$ | $N_r^2(3.418, 0.576)$ |

For the forward fixation when image planes are parallel and coplanar, the horopter was found to be a straight line. In this eye-position, the image planes were at the same distance from the fixation point. Are horopter curves still straight-line horopters in an eccentric gaze with the two parallel image planes, just at different distances from their fixation point? I test this for two fixation points obtained by rotating the fixation point *F*^0^(0, *d*_3.3_). The first fixation point *F*^3^ is the result of rotations by angles *φ_l_* = *φ_r_* = 9.5° and the second fixation point *F*^4^ is obtained by the rotations *φ_l_* = *φ_r_* = −14°.

From the definition of the version angle, *ω* = 1/2(*φ_l_* + *φ_r_*), we see that the fixation points *F*^1^ and *F*^3^ have the same version angle of 9.5°, while the fixation points *F*^2^ and *F*^4^ have the same version angle of −14°. Because the version angle defines the cyclopean axis [7, 26], the fixation points with the same version angle have the same cyclopean axis.

On the image planes for both fixation points *F*^3^ and *F*^4^, in a similar way as was done before for fixation points *F*^1^ and *F*^2^, I take the pairs of corresponding points of the distance *c* = 0, *c* = 0.075 cm, and *c* = 0.125 cm from the points *O_l_* and *O_r_*. The same algebraic method used before to calculate the intersections of the visual lines of the corresponding points now gives the horopter points shown in Table 2.

**Table 2.** The Horopter Points for Fixations *F*^3^ and *F*^4^

| $F^3 : \phi_l = \phi_r = 9.5^\circ$ | $F^4 : \phi_l = \phi_r = -14^\circ$ |
| --- | --- |
| Horopter Points |  |
| $F^3(16.236, 96.916)$ | $F^4(23.419, 93.821)$ |
| $P_1^3(-27.713, 92.943)$ | $P_1^4(12.101, 99.702)$ |
| $Q_1^3(-4.259, 100.973)$ | $Q_1^4(34.016, 88.121)$ |
| $P_2^3(-35.086, 90.340)$ | $P_2^4(4.156, 103.722)$ |
| $Q_2^3(4.003, 103.724)$ | $Q_2^4(40.680, 84.420)$ |

### B. Simulated Horopter Curves

Like the calculations performed above that involve trigonometric functions, simulations in GeoGebra round to 10 decimal places to reach precise geometric conclusions. Numbers are shown here to only their third or fourth decimal place. Moreover, although general conics are fully specified by five points, I used more points to ensure that what I have is the horopter conic and not just a local approximation. The results of the simulations are shown in Figure 7.

**Fig. 7.**
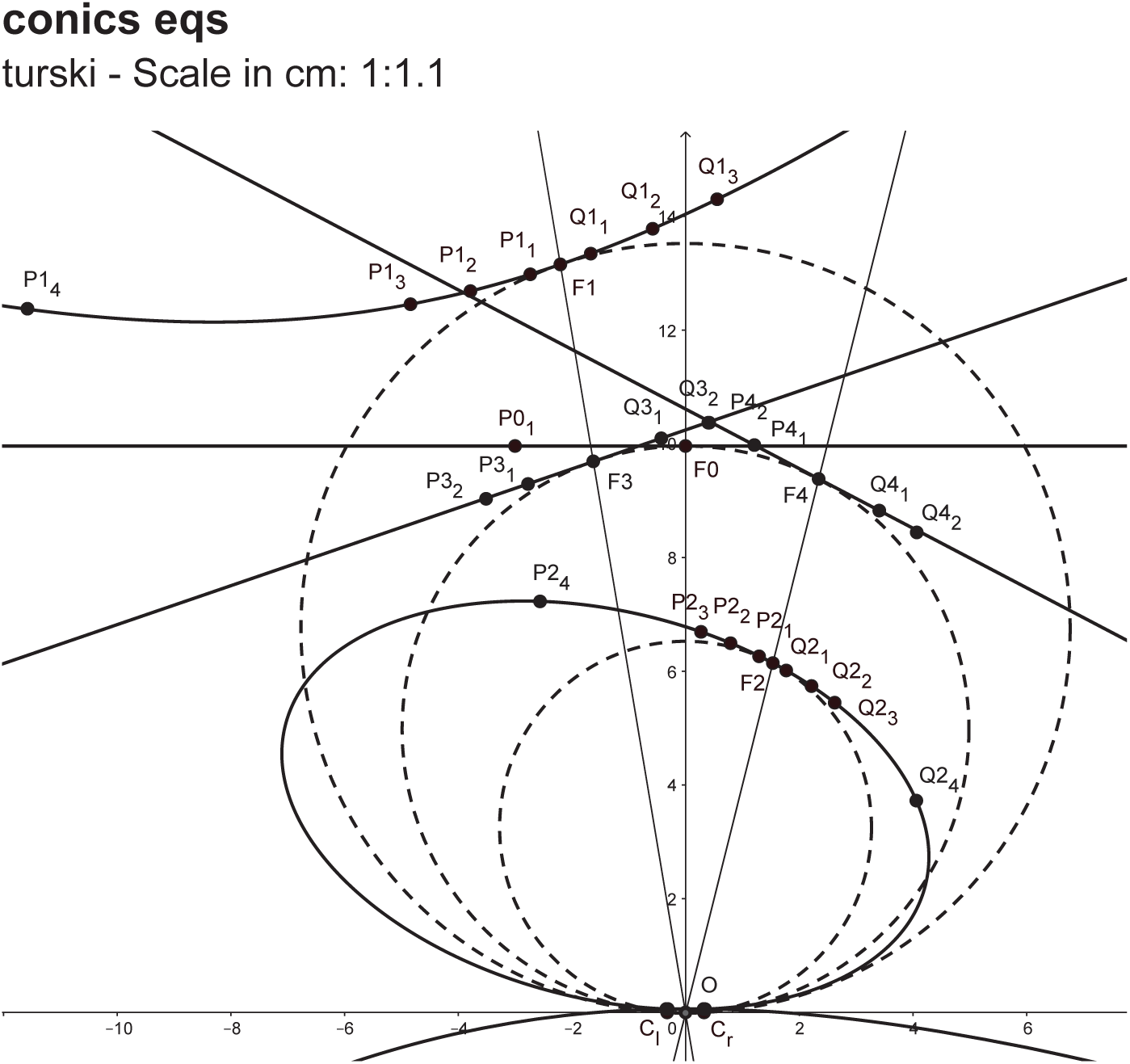
The GeoGebra simulation of five horopter curves for the fixation points: *F*^0^, *F*^1^, *F*^2^, *F*^3^, and *F*^4^. The symmetric fixation *F*^0^ is at the abathic distance for *β* = 3.3°. For the fixation points *F*^0^, *F*^3^, *F*^4^, the image planes of both eyes are parallel, and the horopter curves are straight lines and tangent to the VMC determined by *F*^0^.The units on the axes are in decimeters

GeoGebra also produces the conics equations. Below, I show the equations that GeoGebra generated for the graphs displayed in Figure 7. The numbers are given in decimeters.

Fixation *F*^0^: straight line

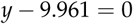

Fixation *F*^1^: hyperbola

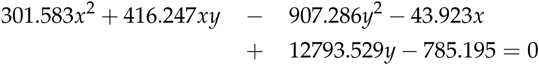

Fixation *F*^2^: ellipse

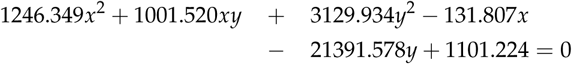

Fixation *F*^3^: straight line

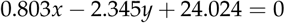

Fixation *F*^4^: straight line

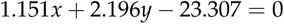

Some of the simulation’s details are missing in Figure 7 due to the scale that is required to display the conic sections’ graphs for the anatomically accurate interocular separation of 6.5 cm and, at the same time, the distances to fixation points, which range from 60 cm to 130 cm. The simulation details near origin *O* of the head coordinate system can be seen in Figure 8, where the viewing window in GeoGebra is magnified 1000 times.

**Fig. 8.**
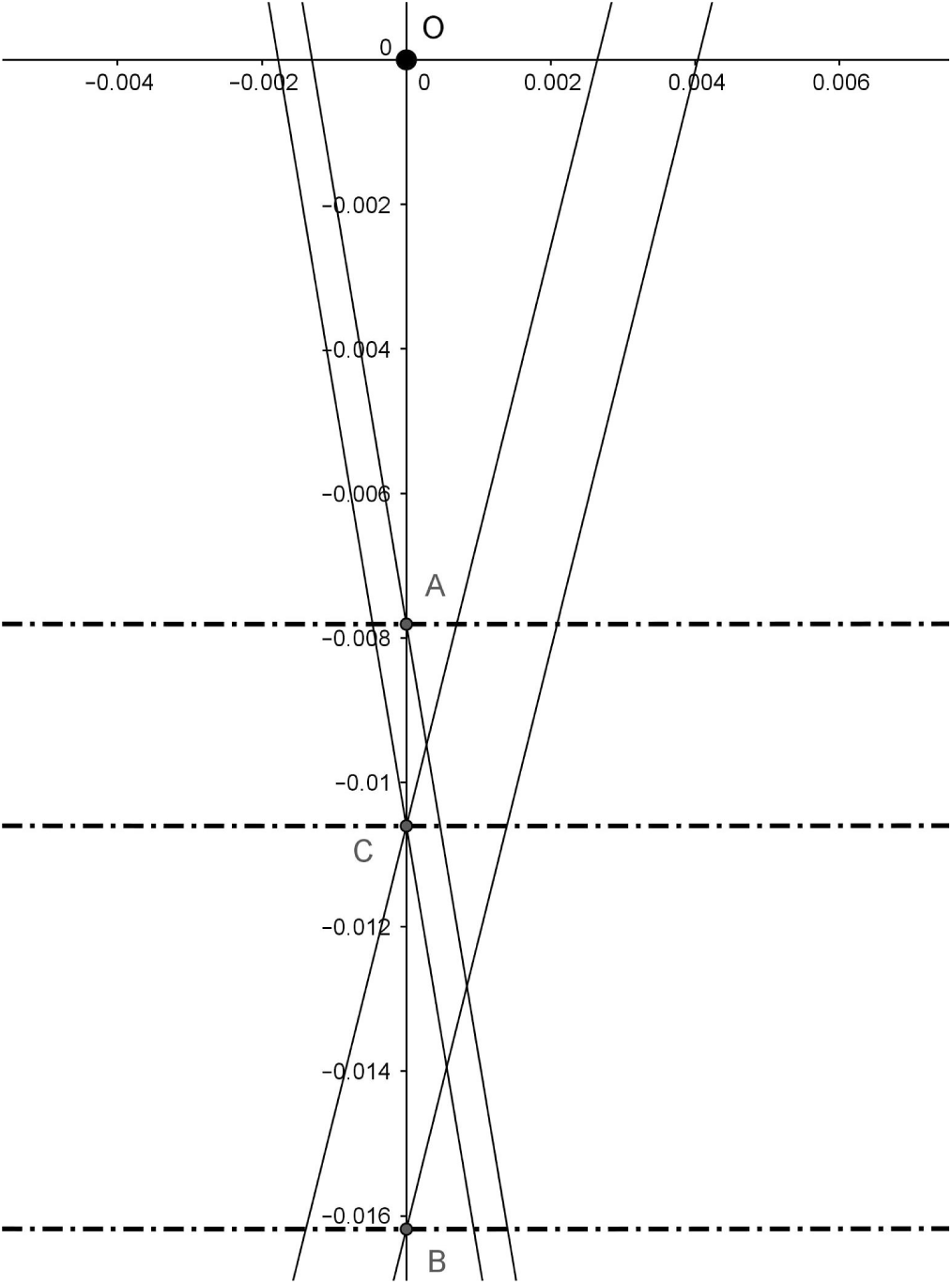
The viewing window near origin O showing the three VMCs in dashed lines. The straight lines through points *A*, *B* and *C* represent the cyclopean axes of the corresponding fixation points with directions specified by the version angles.

The three VMCs through the fixation points *F*^1^, *F*^2^ and *F*^0^, shown in Figure 7 with dashed lines, contain points *A*, *B* and *C*, respectively, though these points are invisible in this figure’s scale. These points are instead displayed in Figure 8 on their corresponding VMCs.

In Figure 8, the vertical line through *A*, *B* and *C* connects to *F*^0^, the symmetrically fixated point at the abathic distance. The two straight lines, one through *A* and the other through *B*, connect to the fixation points *F*^1^ and *F*^2^ respectively. The line with a negative slope that passes through *C* connects to *F*^3^ and the line with a positive slope connects to *F*^4^. These lines all represent the cyclopean axes of their corresponding fixation points. Further, the parallel cyclopean axes have the same version angle value defined with respect to the *y*-axis (the vertical line); lines oriented to the left have a 9.5-degree version and lines oriented to the right have a −14-degree version.

## 5. COMPARISON WITH OGLE/AMIGO CONICS

Using conic equations obtained in GeoGebra for the hyperbola and ellipse, I compute the symmetry axes orientation angles via (11).

The results are the following: for the hyperbola,

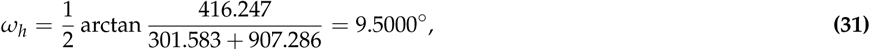

and for the ellipse,

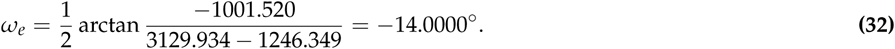

Because of the high level of precision afforded by rounding to ten decimal places in my GeoGebra simulations, I am able to conclude that each conic’s orientation is exactly given by the corresponding version angle. This means that, for a given fixation point, the conic orientation represents the cyclopean axis of the fixation point. Thus, for each fixation point, and as specified by the version angle, the cyclopean axis is the line passing through two points: the fixation point and the point midway between the eyes on the corresponding VMC, cf. [7].

In the Ogle/Amigo approach, for a given asymmetrically fixated point with the azimuthal angle *γ*, the orientation of the conic (6) is given by the angle *ω_c_* in (12). Using the configuration of coordinate systems in Figure 9, I derive the values of *γ* and *b* that are needed to get *ω_c_*.

**Fig. 9.**
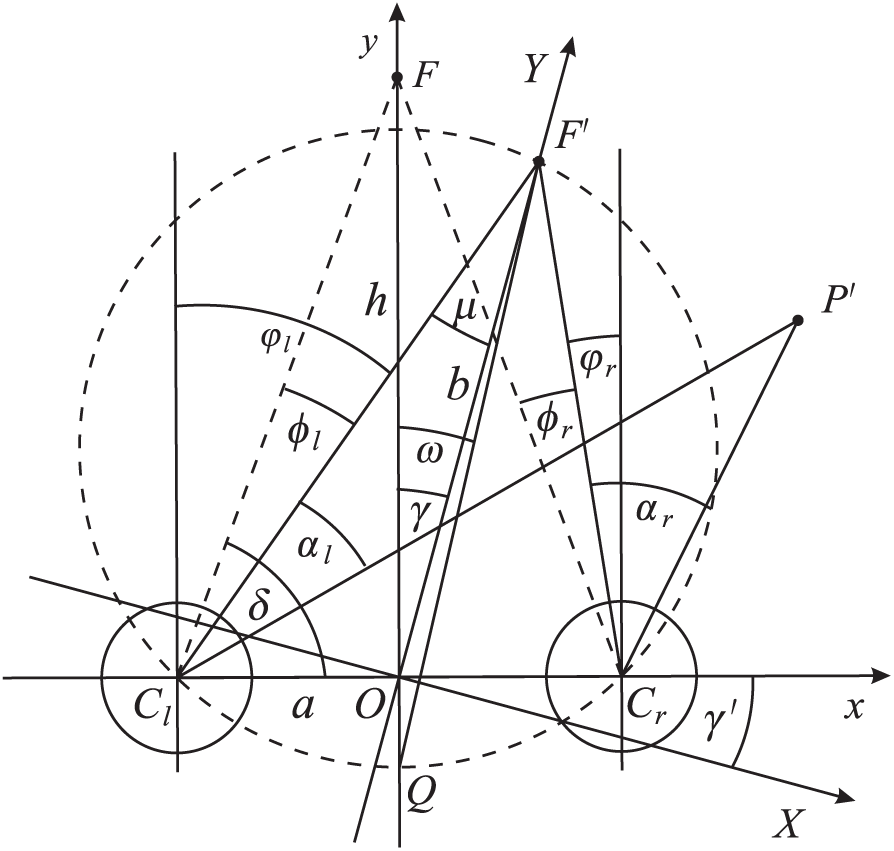
Coordinate systems from the Ogle and Amigo studies are shown with the angles used in calculations comparing results obtained in their studies with those obtained here.

To do this, I first recall the standard expression

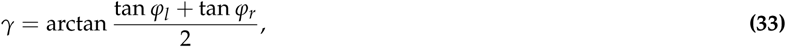

where *ϕ_l_* and *ϕ_r_* are the left and right azimuthal angles shown in Figure 9. The expression in (33) can be easily obtained, for example, from the expressions (9) in [26].

To compute *γ*, I rewrite (33), using simple geometry in Figure 9, as follows:

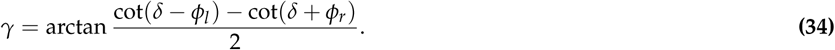

Further, I take *φ_l_* = −15°, *φ_r_* = −13°, *a* = 3.25 cm and *h* = *d*_3.3_ = 99.61 cm, all of which correspond to the fixation point *F*^2^ in our simulation with GeoGebra, and obtain

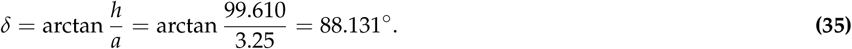

Then, substituting (35) and the above given values of *φ_l_* and *φ_r_* into (34), I compute the angle *γ* as follows:

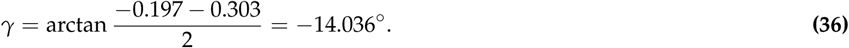

Next, I calculate

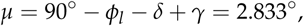

and then, from the Law of Sines,

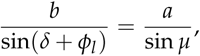

I obtain

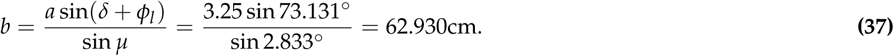

Substituting (36), (37) and *a* = 3.25 cm into (12), I obtain

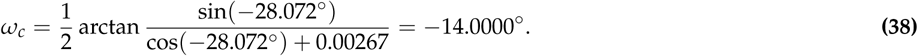

This result in (38) demonstrates that, for the same gaze given by *φ_l_* = −15°, *φ_r_* = −13°, the orientation of the conic in the Ogle/Amigo method is the same as the orientation of the conic obtained in the simulation that is based upon the asymmetric eye model. Moreover, in both cases, the orientation is given precisely by the version angle.

Finally, I explicate the difference between the abathic-distance line horopter given by *H* in (10) and the abathic-distance line horopter simulated using the asymmetric eye. By the Amigo formula (10), the asymmetric fixation *F*′ that is specified by the azimuthal angle *γ* is at the abathic distance given by *b* cos *γ*. Here *b* is the abathic distance to the symmetric fixation *F*^0^ when *γ* = 0.

I prove that *F*′ is on the circle through *O*, *F*′ and *F*^0^, as shown in Figure 10. From the definition of cosine we have the right triangle △*OF*^0^ *F*′ with the right angle at the vertex *F*′. Then, by the well-known result that an angle inscribed in a semicircle is a right angle, *F*′ must be on this circle. In my simulation, the abathic-distance fixation point *F* is on the VMC, shown in Figure 10 by a dashed-line that is passing through *F*^0^ and *O_C_*. We see that, although the numerical values between both cases do not differ significantly, their geometries are indeed quite different.

**Fig. 10.**
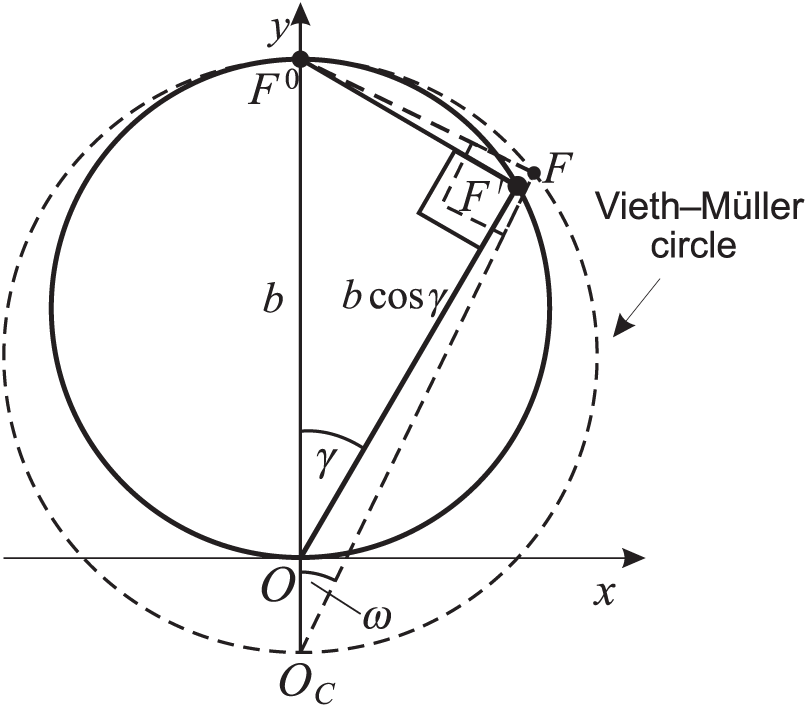
According to Amigo’s formula (10), the fixation point *F*′ with azimuthal angle *γ* is at the abathic distance |*OF*^0^| = *b* cos *γ*. For *γ* = 0, this point becomes the symmetrically fixated point *F*^0^ at the abathic distance, |*OF*^0^| = *b*. I prove in the text that *F*^0^ must be on the circle passing through *O*, *F*′ and *F*^0^. According to my simulation, the abathic-distance fixation point *F* of the version angle *ω* is on the indicated VMC.

## 6. DISCUSSION

The small differences in the perspective projections on the right and left 2D retinae result from the eyes’ lateral separation. The human brain uses this two-dimensional disparity to provide our perception with a vivid impression of depth. The horopter is the locus of all spatial points such that each point is seen in a single direction despite the fact it is projected to the two retinae along different visual lines. When a point lies in front or behind the horopter curve containing the fixation point, the difference in the angles subtended on each retina between the image and the center of the fovea defines absolute retinal disparity. This difference provides a cue for the depth of the object from an observer’s current point of fixation. The difference of retinal disparities for a pair of points defines their relative disparity. This relative disparity provides a cue for the perception of 3D structure such as relative depth and shape.

Here I studied longitudinal horopters in the binocular system with the asymmetric eyes. To render the results of this study in a broader context, I will discuss three eye models of successively increasing anatomical fidelity and the horopter curves that correspond to the binocular system of each model. I will also comment on the prevailing in the literature misinterpretations of the geometric horopter.

The first model is a special case of the reduced eye shown in Figure 4 (a) in which the nodal point is taken to coincide with the eye’s rotation center. The resulting longitudinal horopters are the VMCs originally proposed almost two centuries ago, each passing through the fixation point and connecting the eyes’ rotation centers. When the fixation point moves along the VMC, the rotation centers do not move and, consequently, the circle remains unchanged and the vergence has a constant value. Also, in this model, relative disparity is independent of the eye position [7].

The second model is the reduced eye shown in Figure 4 (a). Here the nodal point is located almost midway between the pupil and the rotation center as required by the eye’s anatomy. I proved in [7] that its longitudinal horopters are circles that pass through the fixation point and connect the nodal points. Further, I proved that, for a given vergence, the circular horopters are parametrized by specific fixation point on the visible part of the VMC and that they intersect at the point of symmetric convergence on this VMC. The vertical component of the horopter is then a straight line passing through this intersection point.

In this model, relative disparity is not independent of eye movement and its changes are within binocular acuity limits. It was conjectured in [7] that these changes in perceived size and shape may not only provide perceptual benefits, such as breaking camouflage, but may also provide the aesthetic benefit of stereopsis [27].

The two reduced eyes, both axially symmetric, show that the VMC differs from the geometric horopter: the VMC connects the eye’s rotation centers whereas the geometric horopter connects the nodal points at their anatomical locations. The main difference is that the relative disparity’s dependence on the eye position applies for the geometric horopters but not VMCs. Although, the VMC provides an acceptable approximation for many applications, the changes in relative disparity may be significant for the finer aspects of depth discrimination, as mentioned above when discussing the second eye model.

The last model, the asymmetric eye, is shown in Figure 4 (b). This is the simplest extension of the reduced eye that includes the foveal displacement from the posterior pole and the cornea and the crystalline lens’s misalignments normally present in human eyes [20, 21]. The longitudinal horopter curves in the binocular system with asymmetric eyes are found, in this paper, to resemble empirical horopters.

Before I continue discussing the impact of the asymmetric eye on the horopter curves, we see that the geometric horopters should be classified into two cases: one for the axially symmetric eye (the second eye model) and the other for the asymmetric eye (the third eye model). And though the VMC provides a useful approximation of the geometric horopters for both symmetric and asymmetric eyes, it should still not be identified with the geometric horopter, as is commonly done in academic education.

In modeling the eye’s asymmetry, I choose to have the fovea’s displacement from the posterior pole be given by an angle *α* of 5.2 degrees. The angle *α* has the least amount of variability among people and it is usually reported in the literature with the value of 5.2 degrees, see for example [22]. This anatomical tilt of the eye relative to the optical axis is the leading cause of aberrations that decrease optical capability. However, the eye’s aplanatic design compensates for these limits through its refractive surfaces’ nonaxial geometry, which includes the cornea and the lens’ misalignment [19].

This study also demonstrates that the asymmetric eye’s refractive surface’ tilt taken in the order of the magnitude of the lens’ misalignment values as measured in healthy eyes produce, in the binocular system with asymmetric eyes, values of abathic distance in the range observed in human vision.

The results obtained here extend the classical studies on modeling empirical horopter curves as conic sections carried out by Ogle [9] and Amigo [10]. The horopter curves derived here are anatomically motivated and have their geometry fully specified by the eye model’s asymmetry parameters, whereas in the studies of Ogle and Amigo, the horopter curves were obtained from an ad hoc introduced equation in [9] and their free parameters had to be determined experimentally for each subject.

In the remaining part of this section, I discuss two problems to which the results obtained here are relevant: vergence resting position and stability in binocular vision.

### A. Vergence Resting Position

During degraded visual conditions, the eyes’ gaze shifts to the resting state of the eye muscle’s natural tonus. This resting position varies across subjects but is relatively stable within a subject. For instance, in darkness, the resting vergence of the eyes, or the dark vergence, ranges from about 30 cm to infinity across subjects [28]. The average value of resting vergence distance across 60 subjects was found in [29] to be 116 cm.

Visual display terminal ergonomic studies have shown that symptoms of visual fatigue during sustained vergence on a target correlate with the distance between the target and the vergence resting position. This correlation has led to our current understanding of resting vergence serving as a zero-reference level for convergence effort [30].

In the binocular system with the asymmetric eye model, abathic distance (14) to the symmetrically fixated points takes on values in the range that is similar to the range of the measured distance of the resting vergence position. Also, both the abathic distance and resting vergence position distance have their average values of about 1 meter. Further, the abathic distance arises in my modeling as the result of the fovea’s anatomical asymmetry, which contributes to the eye optics’ aberrations and the physiological tilt of the crystalline lens that tends, over time, to compensate for various types of aberration [19]. Thus, the natural question that arises is whether or not the abathic distance is closely related, even identical, to the resting vergence distance.

### B. Stability of Binocular Vision

I have proposed here that the principal visual direction for the eye’s position, or the cyclopean axis, is specified by the orientation of the corresponding conic section that models the longitudinal horopter. This specification implies that the principal visual direction is given by the version angle.

As I pointed out in Section 1. A, the perceived directions of all of the points in the visual field are seen in relation to the perceived direction of the point that is being fixated on, that is, the cyclopean axis. However, the cyclopean axis changes direction during eye movements. This means that the cyclopean (fused) image undergoes transformations corresponding to these changes in direction.

Binocular geometry and physiological eye movements cannot be separated from one another; the left and right eyes’ disparate 2D retinal half-images, inherently unstable due to incessant saccadic and smooth pursuit eye movements [31], are fused into a stable 3D representation of the world in the transformation from retinal to cyclopean representations [32].

The outputs from the stereo systems of robotic eyes on a platform replicating saccadic and smooth pursuit eye movements require efficient algorithms for maintaining perceptual stability [33, 34]. The results in this study will be useful to this end.

## ACKNOWLEDGMENT

I thank Michael Landy for reading an earlier version of this manuscript. His helpful comments improved the paper and directed me to the research on vergence resting position.

